# Bioengineered visible polymeric mesh to enhance urogynaecological health

**DOI:** 10.1101/2025.01.23.634411

**Authors:** S Houshyar, A Quigley, I Cole, H. Yin, R Zizhou, T Saha, E Pirogova, JM Yeung, M Mohsenipour, A Mirabedini, E Subejano, AE Shindler, JL Wood, AE Franks, EL Hill-Yardin

## Abstract

Pelvic floor disorders, including pelvic organ prolapse and stress urinary incontinence, are prevalent health concerns, affecting approximately 50% of females over their lifetime, with about 75% of women over 65 years of age being impacted. Traditional surgical interventions, such as transvaginal mesh implants, have led to numerous complications, resulting in their prohibition in several countries. This study introduces an innovative composite mesh designed to mitigate these issues by combining polymethylmethacrylate and thermoplastic polyurethane, further enhanced with iodine-doped carbon nanoparticles to enable visibility via medical imaging. The mesh is coated with 2-methacryloyloxyethyl phosphorylcholine polymer to prevent protein adsorption and promote tissue regeneration. *In vitro* studies showed high cell viability and low protein adsorption, indicating excellent biocompatibility. Implantation of mesh (with or without iodine) in mice revealed no adverse effects on overall animal health. Mouse spleen weight (an indicator of inflammation) was similar between groups; however, levels of some cytokines (i.e., IL-10, IL-17A and GM-CSF) were elevated following implantation of iodinated mesh in mice suggesting that further refinement of the composite mesh is required. Analysis of the fecal microbiome, which is correlated with physiological states, showed that sham and iodinated mesh implant groups maintained consistent microbial profiles with stable diversity (richness and evenness) measures over time. In contrast, the non-iodinated mesh group exhibited decreased species richness post mesh implantation, likely due to a distinct starting microbiome composition prior to implantation. This research is envisaged to contribute to a safer and more effective solution for treating pelvic floor disorders, providing non-invasive post-implantation monitoring and enhanced mechanical compatibility of surgical mesh with native tissue. Our findings demonstrate that this composite mesh possesses mechanical properties that closely mimic human tissue, ensuring biocompatibility, strength, and flexibility without stimulating significant inflammatory or foreign body responses.

## Introduction

Pelvic floor disorders (PFD), including pelvic organ prolapse (POP) and stress urinary incontinence (SUI), present significant health challenges affecting a substantial proportion of adult women worldwide. Epidemiological studies estimate that approximately 50% of women experience some form of PFD during their lifetime, with prevalence rising to about 75% among women aged 65 and older ^1–3^. POP occurs when pelvic organs, such as the bladder, uterus, or rectum, descend into the vaginal canal due to weakened pelvic support structures, including the vaginal walls and surrounding connective tissues ^2–5^. This condition often leads to discomfort, urinary dysfunction, and a reduced quality of life for affected individuals.

Surgical intervention is typically required for managing moderate to severe POP cases. Traditionally, these procedures involved native tissue reinforcement through suturing techniques. However, due to high recurrence rates, transvaginal mesh implants emerged as a more durable solution for patients with advanced POP stages ^1, 6, 7^. While these implants initially showed promise, long-term complications, including chronic pain, infection, erosion, and urinary issues, have emerged in a significant number of cases ^8–10^. Complete mesh removal is particularly challenging, with preoperative imaging and planning being essential. Visualisation of mesh can be complex using radiology ^5, 11^. Concerns over these complications prompted regulatory actions in several countries and led to bans on mesh usage in POP treatment ^12^.

Recent studies have highlighted the critical importance of matching the mechanical properties of implant materials with those of human tissue to reduce complications such as tissue erosion and foreign body reactions. Furthermore, tissue regeneration typically occurs under mechanical stress, promoting a healing response. In the absence of such stress, the surrounding vaginal tissue may be unable to regenerate and instead will degrade over time, increasing the severity of POP symptoms and the risk of mesh erosion and associated complications ^10, 12, 13^. A more flexible mesh can adhere to and more closely match the properties of the underlying tissue ^14^. The mechanical properties of both the mesh and the tissue influence stress transmission at the tissue-implant interface ^14, 15^. Mesh implants need to be optimised to possess similar biomechanical properties to human tissues, facilitating the transfer of applied loads to the surrounding environment to support tissue healing and regeneration ^15, 16^. Although the transvaginal mesh needs to maintain its structure under tension, it must also allow for adaptability and movement during flexion.

As reported in the literature, commercial mesh such as Prolene^TM^, commonly used for POP treatment, is significantly stiffer and stronger than the healthy pre- and post-menopausal vaginal wall tissue (Table 1) ^1, 14, 17, 18^. Therefore, there is a pressing need for innovative biomaterials that replicate the biomechanical properties of human tissue. However, these materials should also offer improved visibility under medical imaging to facilitate monitoring and potential removal post-implantation ^1, 6, 11, 19, 20^. In response to these challenges, we introduce a novel combination of polymethylmethacrylate (PMMA), and thermoplastic polyurethane (TPU) designed to address the critical shortcomings of current treatment options for POP. The composite mesh combines advanced biomaterials with specific features aimed at enhancing clinical outcomes. It incorporates materials that mimic the biomechanical properties of native tissue to support prolapsed organs effectively without causing stress-shielding effects or compromising tissue integrity (Table 1) ^15, 20–22^.

**Table 1.** Comparison of mechanical properties of vaginal wall tissue and commercial polypropylene (Prolene ^TM^, commercially used as TVM)

|  | Post-menopausal vaginal wall <sup>6, 27</sup> | Pre-menopausal vaginal wall <sup>1, 6, 11</sup> | Prolene <sup>TM</sup> <sup>28 29 30</sup> |
| --- | --- | --- | --- |
| **Max Strain (%)** | 1.2 ± 0.05 | 0.8 ± 0.1 | 10 - 12 |
| **Young's Modulus (MPa)** | 12.1 ± 1.10 | 6.7 ± 1.5 | 1350 - 1800 |
| **Max Stress (MPa)** | 0.27 ± 0.03 | 1.7 ± 0.1 | 30 - 35 |

To address the current lack of visibility of implants post-surgery, the use of contrast agents like gold nanoparticles, iodine, gadolinium, and bismuth, which have high K-edge energy but suffer from low solubility and toxicity, are being investigated ^23–25^. Biodegradable carbon nanoparticles, however, offer a promising alternative due to their biocompatibility, fluorescence, and ease of surface modification for bioimaging applications ^11, 23, 24^. Recent studies have explored doping carbon nanoparticles with metallic and non-metallic ions to improve their photoluminescence and X-ray attenuation properties for targeted CT imaging ^15, 20, 23, 26^. With the aim of advancing CT imaging capabilities, the current study specifically investigates iodine-doped carbon nanoparticles that are designed to be non-toxic in physiological environments ^15, 19, 20, 23^.

Furthermore, the composite mesh integrates non-adhesive properties to reduce the risk of unwanted protein adsorption and inflammatory responses- common complications associated with current mesh implants ^31^. These biomaterials, particularly zwitterionic polymers, offer favourable characteristics for permanent and semi-permanent implants. They effectively prevent nonspecific protein adsorption while supporting normal cell growth, regeneration and proliferation in wounded areas ^31^. Among zwitterionic polymers, 2-methacryloyloxyethyl phosphorylcholine polymer (PMPC) has been successfully applied as a coating on various medical devices, including cardiac stents, ventricular systems, vascular prostheses, orthopaedic implants, contact lenses,^32^ and surgical mesh ^31^. Studies demonstrated that PMPC possesses non-fouling, non-toxic, and highly hydrophilic properties both *in vitro* and *in vivo*, making it an excellent choice for enhancing resistance to protein adsorption and promoting biocompatibility, especially for permanent implants ^31–33^.

Herein, a new composite mesh has been developed with several key features, including visibility under medical imaging, biomechanical properties comparable to native human tissue and anti-adhesive properties designed to reduce inflammation by minimising unwanted protein absorption. The newly developed composite demonstrates high biocompatibility in both cell assays and with biological tissue. This mesh provides sufficient strength to support damaged areas without inducing stress-shielding effects on the native tissue. Importantly, it offers excellent visibility under medical imaging, poses no toxicity risk to human tissue and minimises protein absorption, thereby potentially reducing inflammation and foreign body response (FBR). This innovative composite holds promise for treating POP and enables straightforward, risk-free monitoring of post-implantation healing processes.

## Experimental

### Fabrication

The fabrication technique for iodinated carbon nanoparticles (ICPs) followed the method outlined in our previous paper ^23^. In summary, phloroglucinol, iodoacetic acid and cysteine were added to methanol and 2-amino-2-methyl-1-propanol. After a clear solution formed, the mixture was transferred to hydrothermal vessels and heated at 195 °C for 7 h. The resulting brown-yellow products were cooled to room temperature and subsequently underwent dialysis and freeze-drying to obtain dry powders. The fabricated ICPs were characterised for their structural properties using a Burker AXS Discover Diffractometer equipped with a Cu Kα radiation source (λ = 1.5418 Å), operating at 40 kV and 35 mA. X-ray data were collected in the μ-2θ locked couple mode over a 2θ range of 5 - 90◦ (**Fig 1S**). Fourier transforms infrared (FTIR) spectra of materials were recorded using a Perkin Elmer Frontiers spectrometer, with an average of 32 scans per sample and a resolution of 16 cm^-1^ in the range of 4000-700 cm^-1^ (**Fig 2S**). Surface charges (zeta potentials) were measured using a Malvern 885 2000 Zetasizer in deionised (DI) water (**Fig 3S**).

**Figure 1.**
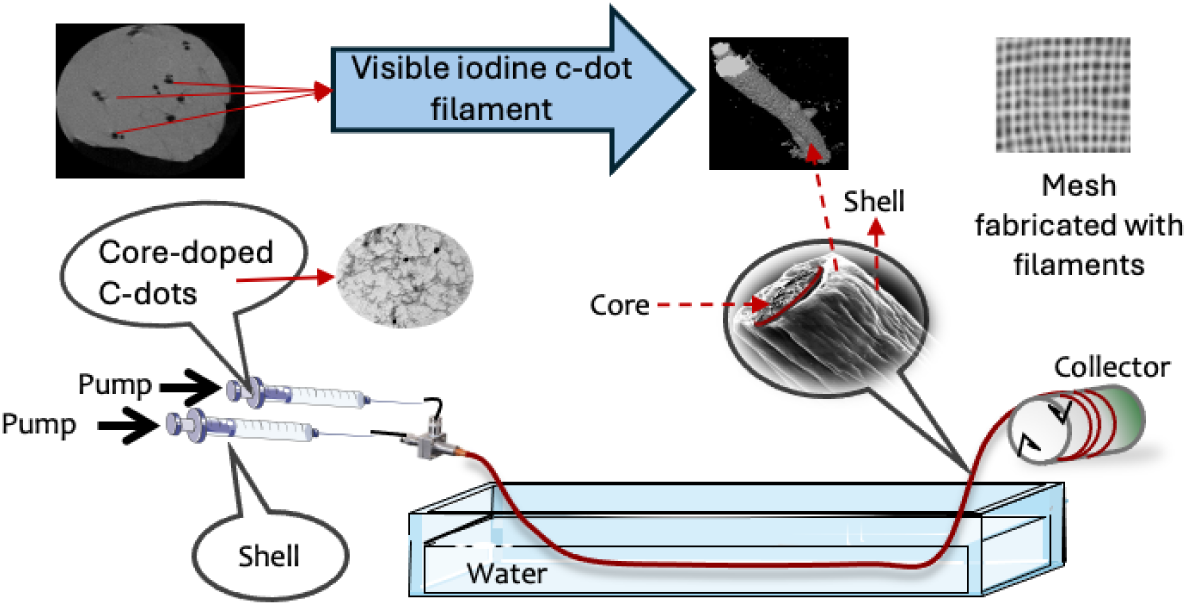
Schematic of the fabrication method of coaxial polymer filament and mesh

**Figure 2.**
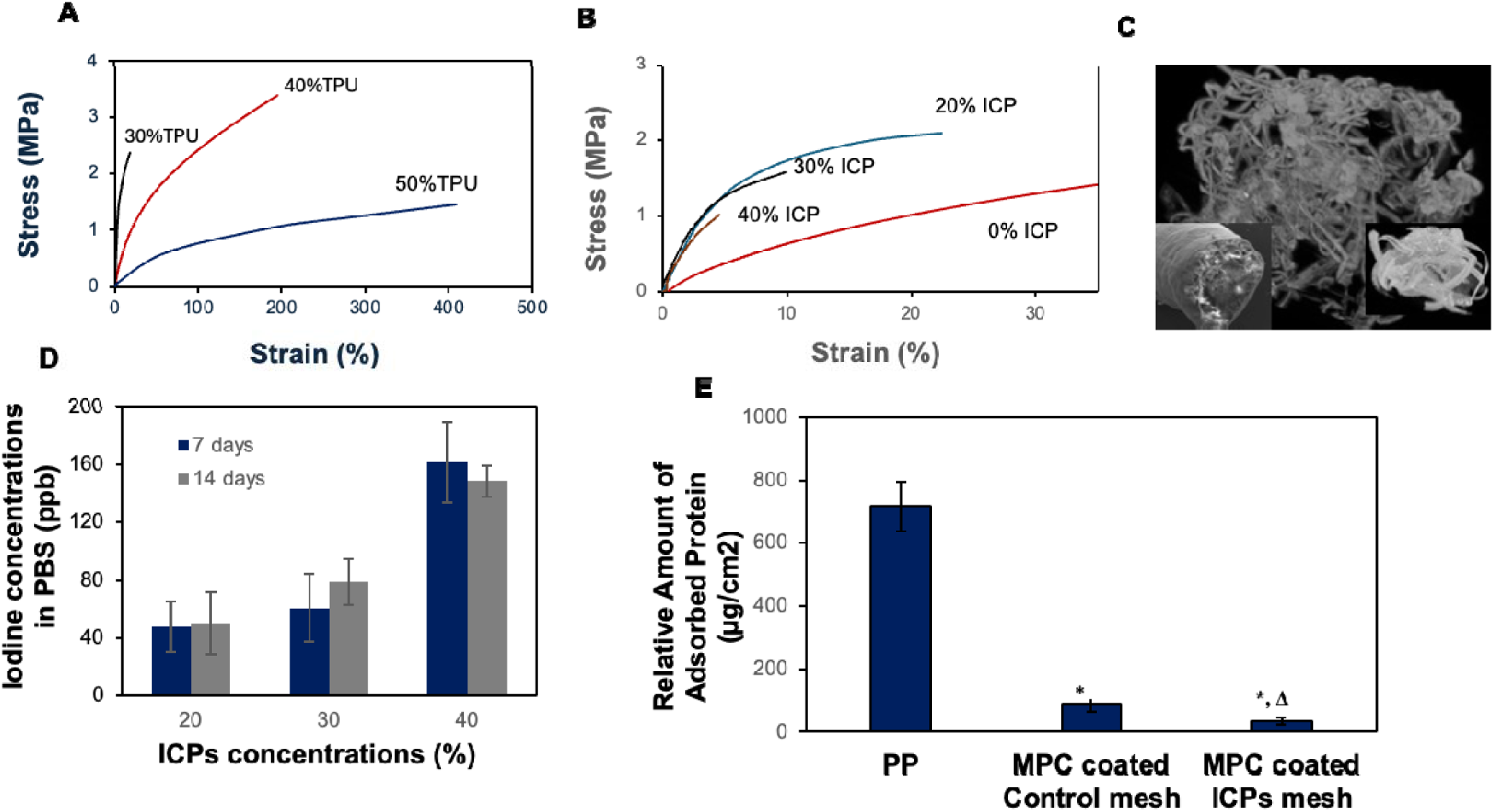
Dynamic mechanical properties of **A:** polymer filaments with various concentrations of PMMA and TPU; **B** filaments with varying ICP concentrations. **C:** MicroCT image of ICP mesh sample. **D**: ICP release rate after 7 and 14 days in PBS. **E:** Protein adsorption levels on PP, PMPC-coated control, and ICP mesh. Symbols *, and Δ indicate a significant difference (n = 3, p < 0.05) in protein adsorption relative to PP and MPC-coated control mesh, respectively, in 2 mg/ml BSA solution.

**Figure 3:**
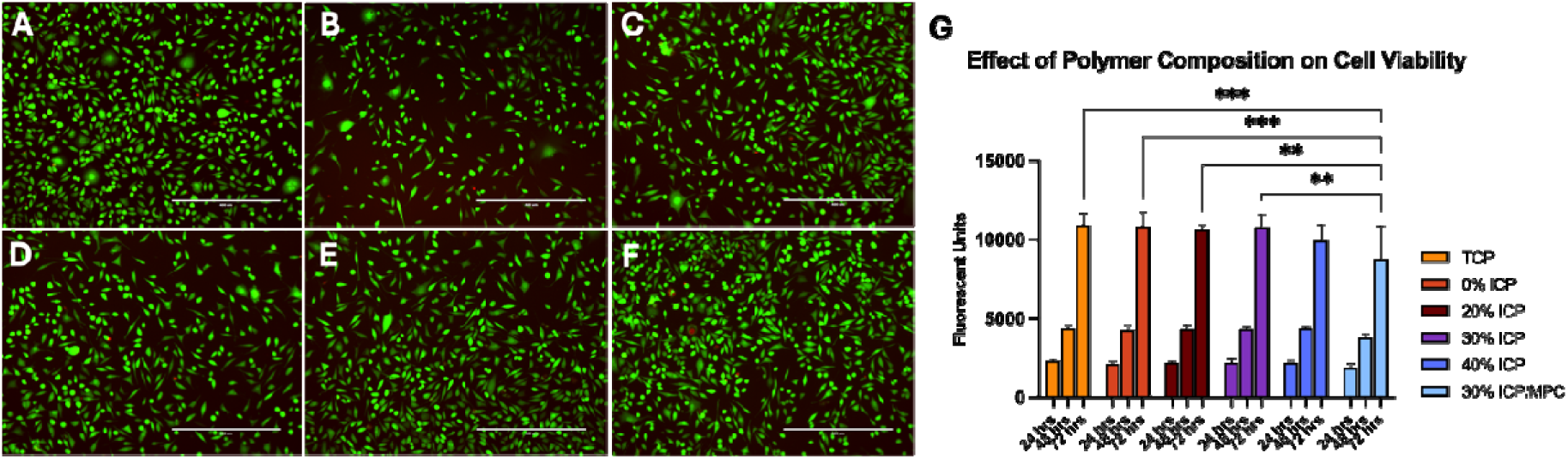
Live/dead viability staining of L929 cells in the presence of polymer samples. L929 cells were grown in the presence of polymer fibres for 72 h to evaluate polymer cytotoxicity. Tissue culture plastic control (A), 0% ICP (B), 20% ICP (C), 30% ICP (D), 40% ICP (E) and PMPC coated 30% ICPs (F). All conditions demonstrated the presence of viable cells at 72 h as detected by live/dead (calcein AM/Ethidium Homodimer) staining (A-F). A quantitative assessment of cell viability was carried out using the cell titre blue assay (G). Polymer formulations, with and without ICP (0-40%) did not show any significant reduction in viability at 24, 48, and 72 h compared to tissue culture plastic (TCP) controls. However, the addition of MPC coating to 30% ICP polymer fibres demonstrated a significant reduction in cell viability at 72 h compared to TCP control and 0% ICP polymers (p<0.001, ***), and 20 % ICP and 30% ICP polymers (p<0.01, **). Scale bars = 400 microns. Error bars represent SD.

The fabrication method for PMMA/TPU filament is illustrated schematically in **Figure 1**. Initially, PMMA was dissolved in DMF at a concentration of 10 wt.% at 50°C for 3h. The prepared PMMA solution was then mixed with a 15 wt.% TPU solution in DMF and stirred for 30 min. The weight ratio of PMMA to TPU was varied at 3:2, 1:1, and 2:3, respectively. ICPs were added to the selected PMMA/TPU ratio of 2:3 based on their biomechanical properties. For the core structure, ICP concentrations ranging from 30% to 60% were incorporated into the polymer solution. The coaxial filament was coagulated in DI water and collected on a drum, with flow rates of 5 ml/hr for the core and 12 ml/hr for the sheath. The collected samples were subsequently dried at 50 °C for several days. After drying, knitted meshes were produced from the coaxial PMMA/TPU filaments using a knitting machine (FAK-Sampler) equipped with a 16-gauge needle.

#### Grafting polymerisation: Coaxial PMMA/TPU (2:3) with 30% and 40% ICPs

After cleaning samples (both control and test) in ethanol for 1 min under sonication, samples underwent grafting with PMPC using UV-irradiated polymerisation techniques. The coating process was undertaken as previously detailed ^31^. In summary, mesh samples were immersed in a benzophenone solution (10 mg/ml in acetone) for 30 s and then dried in the dark at room temperature. Coated samples were subsequently placed in a 0.3 M MPC monomer solution (after removing inhibitors by passing through columns) at a 1:45 mol:lit ratio and exposed to UV radiation (12.78 mW/ cm^2^) for 90 min under an N_2_ gas atmosphere. Finally, PMPC grafted mesh samples were obtained following a 5 min immersion and repeated washing cycles for 10, 5, and 5 min in DI water in a sonicating bath, followed by air drying.

#### Characterisation

Microstructural and elemental analyses were performed using an FEI Verios 460L XHR-SEM equipped with Energy-dispersive X-ray spectroscopy (EDS) @10 kV on a 10 nm iridium-coated filament. Contrast properties were recorded using a Bruker Skyscan 1275 Micro CT @30 kV, with no filter, 360° rotation, and a scanning rate of 2 frames per sec. Attenuated Total Reflectance Fourier Transform Infrared (ATR-FTIR) spectroscopy was used to identify the chemical bonds and functional groups within the composite fibres. FTIR spectra were collected using a PerkinElmer Spectrum-400 spectrometer with 32 scans per spectrum. X-ray photoelectron spectroscopy (XPS) was conducted using a Thermo-Fisher K-Alpha instrument with an Al Kα source. Samples were scanned to a depth of 2–5 nm of the sample surface. Survey spectra were collected with a pass energy of 200 eV, a step size of 1 eV, and dwell time of 50 ms. High-resolution spectra of C1s and I3d were acquired with a pass energy of 50 eV and a step size of 0.1 eV.

Prepared samples were imaged using a Bruker Skyscan 1275 Micro-CT with the previously mentioned imaging parameters. The slices were post-processed into 3D reconstructions using cone-beam reconstruction software (Skyscan NRecon). Image analysis and segmentation were performed using filter and adaptive thresholding with CTAn and CTVol software for quantitative and visualisation purposes. The dynamic mechanical properties of PMMA/TPU, with and without ICPs, were evaluated using a DMA Q800 at 37°C. Samples were tested in universal extension mode (UXF) with a fixed 1Hz frequency and a preloaded force of 0.001N.

The release of ICPs into the media was assessed by immersing the sutures in 5 ml of PBS (pH 7.3) at 37 ^◦^C with shaking at 100 rpm. At various time intervals over 14 days, aliquots were taken for analysis, and the vial was refilled with an equal volume of fresh PBS. The collected liquid was analysed using a UV–Visible-NIR detector (Agilent Carry 5000) across a wavelength range of 200 -800 nm to determine ICP concentration. Quantification of ICPs concentration in the liquid was performed using an Agilent HP 7700 series Inductively Coupled Plasma Mass Spectrometer (ICP-MS). An ICP standard solution was prepared from 99.99% potassium iodide (Chem Supply Australia) within a concentration range of 0 to 100 μg/l. Additionally, the collected filaments after 14 days were analysed using FTIR, EDX, SEM, and XPS, following the previously described methods. The experiments were conducted in triplicate, and the results were reported as an average of the measurements.

To investigate the biocompatibility of the implantable material, a protein adsorption assay was performed following the protocol described in our previous works ^34, 35^. In brief, mesh samples measuring 1 cm × 1 cm were immersed in DPBS at room temperature for 1 h before being incubated in a 2 mg/m1 BSA solution for 2 h at 37 °C. After removing loosely adhered proteins through a series of washes using DPBS buffer solution and Milli-Q water, the adsorbed proteins were desorbed from the mesh surface by incubating the samples at 37 °C in a 1% w/v sodium dodecyl sulphate solution for 1 h. A protein quantification kit utilising Coomassie Brilliant Blue G (CBB) dye was employed to interact with both standard and test (desorbed) protein solutions, producing a blue colouration. The intensity of the colouration was measured in terms of optical density (OD) values (n = 3) using a SpectraMax Paradigm Multi-Mode Detection Platform (Molecular Devices, USA). The concentrations of test protein solutions (desorbed from mesh) were calculated from their OD values of the test solutions and the slope of the standard curve, which was plotted with the concentration of standard protein solutions (X-axis) and their corresponding OD values (Y-axis). Mean values were compared using a Student’s t-test (Microsoft Excel) with p values less than 0.05 considered to indicate statistical significance.

### *In vitro* cytocompatibility

Polymer fibres were cut into 5 mm lengths and placed in the wells of a 96-well plate (Greiner) in quadruplicate. Samples were sterilised with 70 % ethanol for 5 min, air-dried, and washed twice with calcium- and magnesium-free PBS (Gibco). L929 cells were then seeded into each well (5,000 cells/well) in growth media (DMEM (Gibco) supplemented with 10% FBS (Gibco), 2 mM L-glutamine (Gibco), and penicillin/Streptomycin (Gibco)). Cells were allowed to attach and proliferate for 24, 48 and 72 h in the presence of polymer samples, followed by cell viability analysis.

L929 cell viability was assessed via live/dead staining at 72 h. As described above, the polymer fibres were sterilised, cut into 5 mm lengths and placed into a 96-well plate. L929 cells were seeded (5,000 cells per well) into wells containing the polymer samples, with tissue culture plastic used as a control. Cells were allowed to attach and proliferate for 72 h before live/dead staining. After 72 h of incubation, to assess live/dead staining, Calcein AM (Thermo) and Ethidium Homodimer (Thermo) were added to each well at final concentrations of 1 mM and 0.5 mM, respectively, in growth media. Cells were incubated under standard tissue culture conditions for 30 min. After incubation, the staining solution was removed and replaced with fresh growth media. Images of each well were immediately captured using an EVOS digital microscope (Thermo).

Metabolic viability was assessed using a cell titre blue assay kit (Promega), according to manufacturer instructions. L929 cells were seeded into the wells of a 96-well plate containing polymer samples, as described above (5,000 cells/well). Cells were allowed to attach for 24, 48 and 72 h before analysis of metabolic activity was carried out at each time point. To analyse metabolic activity, the cell assay solution was prepared by mixing cell titre blue reagent with growth media at a ratio of 1:5. The culture media was replaced with the cell assay solution, and the assay was allowed to proceed for 3 h before absorbance was assessed using a fluorescent plate reader (CLARIOstar, BMG Labtech). Statistical analysis (2-way ANOVA with Tukey posthoc analysis) was carried out using GraphPad Prism (version 10.2.2).

### In vivo study

#### Animals

Twelve six-week-old female BALB/c mice were purchased from the Australian Resource Centre (ABR, Sydney, Australia) and used for experiments. Mice were housed in 3 cages of 4 mice, prior to surgery. Mice were acclimatised at the RMIT University animal facility for 2 days (Day 0 and Day 1) prior to commencing faecal sample collection on Day 2 (pre-surgery), Day 8 (one day post-surgery), and Day 14 (day of cull). On Day 2 mice were placed in individual alcohol-cleaned plastic container for a maximum of 1 h to obtain a faecal sample and then returned to the original cage. Mice were housed together during acclimatization, then singly housed for 5 days from day of surgery to enable individual recovery, then on Day 11 mice were returned to previous cages. On Day 12 mice were placed into 2 cages (containing 5 and 6 mice from sham and mesh-implanted groups).

#### Faecal sample collection

Faecal samples from control (sham), iodine mesh-implanted, and non-iodine mesh-implanted groups were collected pre-surgery (Days 2, 3, 4, 5, and 6), post-surgery (Days 8, 9, 11, 12, and 13), and at cull (Day 14) for gut microbiome analysis, yielding 125 samples. All samples were immediately stored at -80°C upon collection to preserve microbial integrity until further processing.

#### Surgical procedure

On day 7, mice underwent surgery to implant a 1 cm x 1 cm mesh sample within the subcutaneous space (under the skin, on top of the muscle-deepest layer of skin) on the rear flank. There were three groups of mice, with four mice in each group. Test group 1 underwent sham surgery (surgery was undertaken but mesh was not implanted). Test group 2 underwent surgery to implant a 1 cm x 1 cm mesh sample without ICPs. Test group 3 underwent surgery to implant a 1 cm x 1 cm mesh sample containing ICPs. Mice were euthanised by intraperitoneal injection of ketamine (100 mg/kg) and xylazine (10mg/kg) as approved by the RMIT Animal Ethics Committee (project #25880), 7 days after surgery.

Anaesthesia was induced using 3-5% isoflurane in an induction chamber. An incision in the dorsal thoracic region of the mice was made with a sterilised scalpel, and the mesh sample was inserted under the skin. Following the surgical procedure, the incision was closed using surgical staples (7 mm wound clip; Able Scientific #AS59035). During surgery, body temperature was maintained at 37 °C using a heating pad (Adloheat N11384), regulated via a rectal temperature probe feedback. After surgery, the mice were continuously monitored while placed on the thermal pad until they began to move independently. Pain relief (5 mg/kg meloxicam and 0.05 mg/kg buprenorphine) was administered via intraperitoneal injection, and the animals were transferred to prewarmed cages with thermal pads. Post-surgery, animals were monitored every 30 min for 2 h, followed by hourly checks for the next 4 h. A second dose of pain relief (5 mg/kg meloxicam and 0.05 mg/kg buprenorphine) was administered via intraperitoneal injection 24 h post-surgery. The mice were single-housed for 4 days post-surgery to prevent injury from the other cohoused mice and monitored once daily after that.

#### Cytokine assay

Blood was collected post-mortem via cardiac puncture to measure inflammatory cytokines. In brief, the cardiac puncture procedure entailed making a lateral incision through the upper abdominal wall beneath the rib cage. The diaphragm and rib cage were dissected towards the collarbone, and the sternum and rib cage were retracted to expose the thoracic cavity. A 25G needle was inserted into the right ventricle, and a 1 ml syringe attached to the needle was gently retracted to obtain 0.5-1.0 ml of blood.

Cytokine levels in cardiac blood were measured using a Bio-Plex Pro Mouse Cytokine Th17 Panel A 6-Plex Immunoassay, together with a Bio-Plex Pro mouse IL-22 immunoassay kit (Bio-Rad 171GA004M, M6000007NY, Bio-Rad, Hercules, CA). The assay was conducted according to the manufacturer’s instructions, and cytokine concentrations were obtained using a Bio-Plex 200 system (BioRad). Measurements followed the BioRad Bio-Plex Pro assay instruction manual (#10014905) for mouse cytokine groups I and II.

#### DNA extraction and 16S rRNA gene sequencing

Bacterial genomic DNA was extracted from faecal samples using the DNeasy PowerSoil Pro Kit (QIAGEN), following the manufacturer’s protocol. The V3-V4 region of the 16S rRNA gene was amplified using primers 341F and 805R under the following PCR conditions: initial denaturation at 98°C for 30 s, followed by 25 cycles of denaturation at 98 °C for 10 s, annealing at 55 °C for 30 s, and extension at 72°C for 30 s, with a final extension at 72°C for 2 min. Dual indexing was performed using Nextera XT Index Kit (Illumina) to allow for library multiplexing. PCR reactions were carried out on a CFX Connect thermal cycler (Bio-Rad). The PCR products were purified using AMPure XP beads (Beckman Coulter) and quantified with the Qubit dsDNA BR Assay Kit (Invitrogen). Purified libraries were pooled in equimolar concentrations and sequenced using paired-end reads (2x300 bp) with the MiSeq Reagent Kit v3 on an Illumina MiSeq System.

#### Sequencing data analysis

The sequencing data were processed and analysed using the QIIME2 pipeline (2024.2 release) ^36^. Quality control and denoising were performed with DADA2 ^37^ , while taxonomic classification was performed using a Naive Bayes classifier trained on the SILVA SSU Ref NR 99 138.1 database ^38^. After classification, microbial community analysis, including alpha and beta diversity metrics, was performed. Visualisation and statistical analysis were performed on RStudio (v2023.06.0+421) using the phyloseq, qiime2R, microbiome, and vegan packages ^39–44^.

## Results and discussion

### Fabrication

The synthesis process yielded a dark brownish product, which was characterised according to the methodology as previously published ^20, 23^. XRD analysis confirmed the presence of ICPs, showing a peak at 19.5°, corresponding to the (002) plane of carbon interlayer spacing (d=0.46 nm), indicative of an amorphous carbon structure ^23^. Additionally, the particles exhibited a porous structure with a surface that displayed a negative charge with a zeta potential of -40 mV, suggesting the presence of functional groups, particularly carboxylic and hydroxylic groups, as shown in **Supplementary** Figure 1. The FTIR spectrum, as depicted in **Supplementary** Figure 2, demonstrated the chemical structure of the ICPs, further revealing absorption peaks corresponding to hydroxyl, amide and carbonyl groups alongside peaks corresponding to iodine (I-O) at 440 cm^-1^ and 580 cm^-1^.

ICPs were incorporated into PMM/TPU filaments using a wet spinning process. To optimise the dynamic mechanical properties using a dynamic mechanical analyser (DMA) of the filaments for surgical mesh applications, three compositions of PMM and TPU were formulated: 2:3, 1:1 and 3:2. **Figure 2A** shows the DMA properties of the filaments. The 50/50 PMM/TPU composition exhibited excessive stretchiness and flexibility at body temperature, combined with low stiffness. Conversely, the 2:3 composition was overly stiff, with limited flexibility. Incorporating ICPs into the materials increased stiffness while reducing flexibility across all compositions. Consequently, the 40/60 TPU/PMMA filament demonstrated the most optimal mechanical properties in terms of flexibility and stiffness, making it suitable for surgical mesh fabrication. As a result, a filament composed of 40 % TPU and 60% PMMA, along with varying concentrations of ICPs, was fabricated using the same wet spinning technique. As expected, DMA results for the materials at body temperature showed that the addition of ICPs increased stiffness while decreasing flexibility **(Figure 2B)**. Samples with ICP concentrations of 20%, 30%, 40% and 60% were produced. However, samples containing 60% ICPs proved too brittle, rendering them unsuitable for further characterisation and production. Among the tested samples, the filament containing 30% ICPs exhibited the highest stress and strain at break, measuring 1.92 MPa and 7.5%, respectively. Further increases in ICP concentration reduced stiffness and flexibility, with the sample containing 40% ICP recording stress at a break of 1.1 MPa and a strain of 4.5%. Based on these findings, filament containing 30% ICPs was deemed optimal for producing surgical mesh, as its mechanical properties closely align with those of pre-menopausal vaginal tissue (1.7 MPa stress at break, 1.1% strain) and were much lower than commercially available surgical mesh (30-35 MPa stress at break, 10-12% strain).

The contrast properties of the filament observed via CT scan (**Figure 2C)** were also quantified. The filament containing 30% ICPs exhibited a contrast value of 382 Hounsfield units (HU), which falls within an acceptable range. For reference, water typically measures 0 HU, adipose tissue ranges from -20 to -150 HU, kidney tissue ranges between 20-50 HU, and dense bone registers around 1000 HU.

To evaluate the release of ICPs into the surrounding environment and assess the potential impact on biological systems and imaging visibility, the release rate of ICPs in PBS was monitored for 7 and 14 days at 37°C with mild agitation (**Figure 2D**). The release rate was measured by quantifying the amount of iodine released from ICPs into the PBS solution using both Inductively Coupled Plasma Mass Spectroscopy (ICP-MS) and UV-Vis spectroscopy. Results indicated that the amount of iodine released into the PBS solution remained minimal for all samples, reaching a plateau after 7 days. This suggests that most of the iodine remained encapsulated within the filament’s core section due to the protection provided by the PMMA/TPU shell. The initial iodine release into the media may be attributed to residual iodine deposited during fabrication or from the exposed ends of the filaments. In-vitro and in-vivo testing were conducted to investigate the potential toxicity of released iodine on tissue (detailed in the cell and in vivo testing section). After 14 days in PBS, the filament was chemically analysed using FTIR. The presence of ICPs within the filament was confirmed by the presence of the C=C stretching vibration peak at 1610 cm^-1^. However, the peak intensity was lower than the virgin filament, likely due to the minimal release of ICPs into the surrounding media (see **Supplementary** Figure 2).

#### Protein adsorption properties

The protein adsorption test result provides a valuable preliminary indication of the immune response elicited by an implant within the host body. Lower protein adsorption on the implant material suggests a reduced immune response, promoting better implant integration with minimal complication. In this work, we coated the filament with poly PMPC to mitigate protein adsorption. PMPC, known for its zwitterionic nature, exhibits no toxicity and demonstrates anti-thrombogenicity, high hydrophilicity, and resistance to bacteria and protein adsorption. Knitted samples from the prepared coated filaments were tested for protein absorption properties. Protein absorption properties of commercial surgical mesh composed of polypropylene (PP), PMPC-coated mesh made of 30% ICP filament and control mesh samples made of 0% ICP filament are shown in **Figure 2E**. Protein adsorption on a substrate is considered a crucial parameter for assessing the biocompatibility properties of implant materials during the early stages of product development. As evident from the graph, both PMPC-coated mesh samples showed a significant reduction in absorbed BSA protein (2 mg/ml BSA protein solution) compared to the uncoated PP mesh. The PMPC-coated control mesh showed an 87.8% reduction in protein adsorption relative to the PP mesh, emphasising the anti-adsorptive nature of PMPC due to the thick hydration layer formed by water molecules surrounding the methyl groups of PMPC^35^. The comparison between the PMP-coated control and the PMPC-coated ICP mesh is of particular interest. The latter exhibited a significantly lower amount of adsorbed protein (n = 3, p < 0.05), suggesting an additional effect of ICPs in reducing protein adsorption on the coated mesh surface. Specifically, protein adhesion for the PMPC-coated ICP mesh was measured at 35.3± 10 µg/cm^2^, representing a significant reduction of 95% and 6% compared to the PP mesh and PMPC-coated control mesh, respectively. These findings suggest enhanced biocompatibility of the PMPC-coated ICP mesh if utilised as an implantable device.

### *In vitro* analysis of cytocompatibility

Cell viability and proliferation were assessed using L929 cells, which are commonly used for the biological evaluation of medical devices ^45^. To assess the effects of ICP loading and PMPC coating on cellular viability and proliferation, L929 cells were seeded into wells containing polymer formulations with varying ICP concentrations, both with and without MCP coatings. Analysis of cell viability *via* live/dead staining revealed over 90% live cells across all conditions tested at 72 h (including tissue culture plastic control (TCP), 0% ICP, 20% ICP, 30% ICP, 40% ICP, and MCP coated 30% ICP fibres, as seen in **Figure 3 (A-F)**, suggesting minimal cytotoxicity of all tested formulations. Cell viability and proliferation were further assessed using the cell titre blue assay after 24, 48 and 72 h of exposure to the polymer fibres. The results indicated no significant toxicity across all polymer formulations, as L929 cell viability was maintained at 24, 48 and 72 h (**Figure 3G**), compared to the tissue culture control samples, with one exception: L929 cells exposed to MPC-coated polymer fibres with 30% ICP for 72 hours exhibited a significant reduction in viability. This suggests that the MCP coating may influence cell proliferation or metabolism. Furthermore, the 72 h exposure of L929 cells to MCP-coated 30% ICP polymer resulted in a significant reduction in cell viability compared to L929 cells exposed to uncoated 30% ICP polymer, as well as those exposed to 20% ICP and 0% ICP formulations under the same conditions.

In summary, the presence of ICP (0 – 40%) did not significantly affect cell viability (or proliferation) in the formulations tested. However, the MPC coating on the 30% ICP-loaded mesh samples significantly reduced L929 cell viability (approximately 19%) after 72 h. Nevertheless, all polymer formulations supported cell proliferation over the 24–72 h period.

### In vivo study

#### Animal health

One animal from the sham group (i.e., underwent surgical procedure without mesh implantation) did not recover from anesthesia (an adverse event report was logged with the relevant animal research committee and no ethical issues were detected). No other mice showed signs of ill health throughout the experimental protocol.

#### Spleen weight

Mouse spleen weights were measured to assess potential systemic effects of the surgically implanted mesh relevant to the overall inflammatory status of the mice. There was no difference in mouse spleen weight between animals in the control (sham), implanted mesh with IPCs or without IPCs groups (**Figure 4A**). The average spleen weight for sham surgery mice was 57.03 ± 1.2 mg (n=3 mice, mean ± sem). Mice implanted with iodine IPC mesh had an average spleen weight of 58.95 ± 2.0 mg (n=4 mice), while mice implanted with non-iodinated mesh (i.e., mesh without IPCs) exhibited an average spleen weight of 64.37 ± 3.6 mg (n=3 mice). There were no statistically significant differences in spleen weight between the three groups (Sham versus iodinated ICP mesh: p = 0.838; sham versus non-iodinated mesh: p = 0.170; iodine ICP mesh versus non-iodinated mesh: p = 0.300). The absence of differences in spleen weight between mice that underwent sham surgery and those that received surgical implants suggests that the procedures did not induce significant changes in systemic inflammatory status.

**Figure 4.**
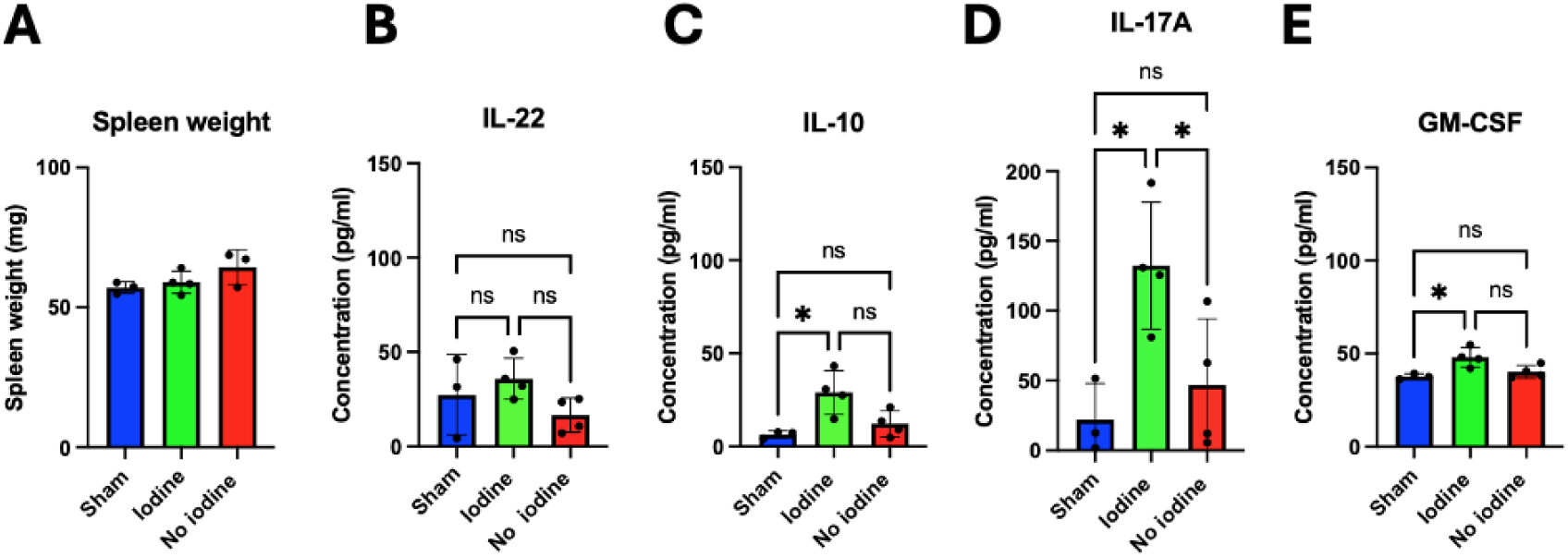
Mouse spleen weight and cytokine levels post-surgical implantation. **(A)** Mouse spleen weight. Spleen weight (mg) from mice that underwent sham surgery (no mesh implant), surgery with a non-iodinated mesh implant (w/o iodine), or surgery with iodinated ICP-containing mesh implant (w iodine). Serum cytokine levels. Serum cytokine levels: **B:** IL-22, **C:** IL-10, **D:** IL-17A **E:** GM-CSF. GM-CSF: granulocyte-macrophage colony-stimulating factor, IL-22, -10,-17A: Interleukin-22, -10, -17A, respectively. One-way ANOVA with Tukey’s multiple comparisons test. *p<0.05.

#### Serum cytokine levels

To assess for peripheral inflammation, we measured major cytokines in serum (IL-22, IL-10, IL-17A and GM-CSF). These cytokines were selected to provide a broad profile of potential inflammatory responses in mice following implantation of mesh with iodine (ICPs) compared to non-iodinated mesh and sham surgery.

#### IL-22

Measurements of mouse serum from mice that underwent sham surgery, mesh implant including iodinated ICPs, and mesh implant without ICPs showed no statistically significant changes in IL-22 cytokine serum levels (**Figure 4B**). Serum samples from sham-operated mice had a mean of 27.37 ± 12.3 pg/ml (n=3 mice) of IL-22. Serum samples from mice that received a mesh implant containing iodinated ICPs had a mean level of 35.86 ± 5.4 pg/ml IL- 22 (n=4 mice). Mice that received an implant of mesh lacking ICPs had a mean level of 16.61 ± 4.5 pg/ml (n=4 mice) of IL-22. There was no difference in the levels of IL-22 in sham versus iodine-implanted mice (p=0.707). Similarly, we observed no differences in IL-22 levels for sham versus implant without iodine (p=0.582). In addition, IL-22 levels for iodine versus without iodine implants were similar (p=0.178).

#### IL-10

Our analyses showed an increase in IL-10 cytokine serum levels in mice implanted with ICP mesh compared to mice that underwent sham surgery. There was no difference in IL-10 levels between ICP mesh-implanted mice and those that received a non-ICP-containing mesh implant (**Figure 4C**). Samples from sham-operated mice (n=3) had a mean of of 6.37 ± 1.1 pg/ml of IL-10. Mice that received a mesh implant containing ICPs (n=4 mice) had a mean level of IL-10 of 29.0 ± 5.9 pg/ml. Mice that received a mesh implant lacking ICPs (n=4 mice) had a mean level of 12.15 ± 3.5 pg/ml of IL-10. IL-10 levels were increased in mice with ICP mesh implants compared to sham surgery (p=0.019). There was no statistically significant change in IL-10 levels between the sham group and mice implanted with mesh without iodine (p = 0.655). There was an increase in IL-10 levels in mice implanted with ICPs compared to mice that received mesh without iodine. However, this did not reach statistical significance (p = 0.052).

#### IL-17A

We observed an increase in IL-17A cytokine serum levels in mice implanted with ICP mesh compared to those that received an implant lacking ICPs (p=0.050). Similarly, mice with ICP implants had higher IL-17A levels compared with mice that underwent sham surgery (p=0.022). A similar level of IL-17A was measured in mice that received the mesh implant without iodine and those that underwent sham surgery (p=0.732) (**Figure 4D**). Sham-operated mice had a mean IL-17A level of 22.00 ± 15.1 pg/ml (n=3 mice). Mice that received a mesh implant containing ICPs had a mean serum IL-17A level of 132.3 ± 22.7 pg/ml (n=4 mice), while mice that received a mesh implant lacking ICPs had a mean serum IL-17A level of 46.78 ± 23.7 pg/ml (n=4 mice).

#### GM-CSF

Serum cytokine data showed increased GM-CSF levels in mice that received a surgical implant containing mesh with ICPs compared to sham-operated mice (p=0.02). There was no difference in GM-CSF levels between sham-operated mice and those that received a non-ICP-containing mesh implant (p=0.678). GM-CSF levels were higher in mice with ICP-containing mesh than those with non-iodinated mesh, but this increase did not reach statistical significance (p=0.052) **(Figure 4E**). Sham-operated mice had a mean GM-CSF level of 37.72 ± 0.8 pg/ml (n=3 mice). Mice that received a mesh implant containing ICPs had a mean serum GM-CSF level of 48.06 ± 2.6 pg/ml (n=4 mice), while those that received a mesh implant lacking ICPs had a mean level of 40.28 ± 1.7 pg/ml (n=4 mice).

Although there were no overall health implications observed during routine monitoring of the mice, these findings suggest that the presence of the iodinated mesh implant triggered an inflammatory response in the host organism. Levels of the anti-inflammatory cytokine, IL-10, and GM-CSF, a cytokine involved in wound healing, were elevated in mice implanted with ICP mesh compared to sham-operated mice. Similarly, production of the proinflammatory cytokine IL-17A was elevated in mice with ICP mesh implants compared to mice that underwent sham surgery. IL-17A levels were also significantly higher in mice with ICP mesh implants compared to those with non-iodinated mesh surgery. Collectively, these findings indicate that, while well tolerated, additional measures to refine the properties of the mesh are required to combat potentially non-beneficial inflammatory responses in the host organism in response to the iodinated mesh implantation.

### Fecal microbiome analysis

#### Relative abundance

Following the analysis of fecal samples collected pre-surgery, immediately post-surgery and at the time of cull, Bacteroidota and Firmicutes were the predominant phyla across all groups of the study **(Figure 5A)**. The fecal microbiome from the sham control and iodine mesh groups appeared more similar over time compared to the no-iodine mesh implanted group. Despite the absence of implanted mesh, control (sham) group samples showed fluctuations in the relative abundance of Verrucomicrobiota, accompanied by an expansion of Proteobacteria. A marginal decrease in the relative abundance of Firmicutes was observed in the iodine mesh group post-surgery, as along with fluctuations in Verrucomicrobiota levels. In the non-iodinated mesh group, we observed an expansion of Bacteroidota, accompanied by a decrease in the proportion of Firmicutes. In contrast with the sham control and ICP (iodinated) mesh groups, the no-iodine mesh group displayed a small proportion of Verrucomicrobiota that remained stable throughout the study.

**Figure 5.**
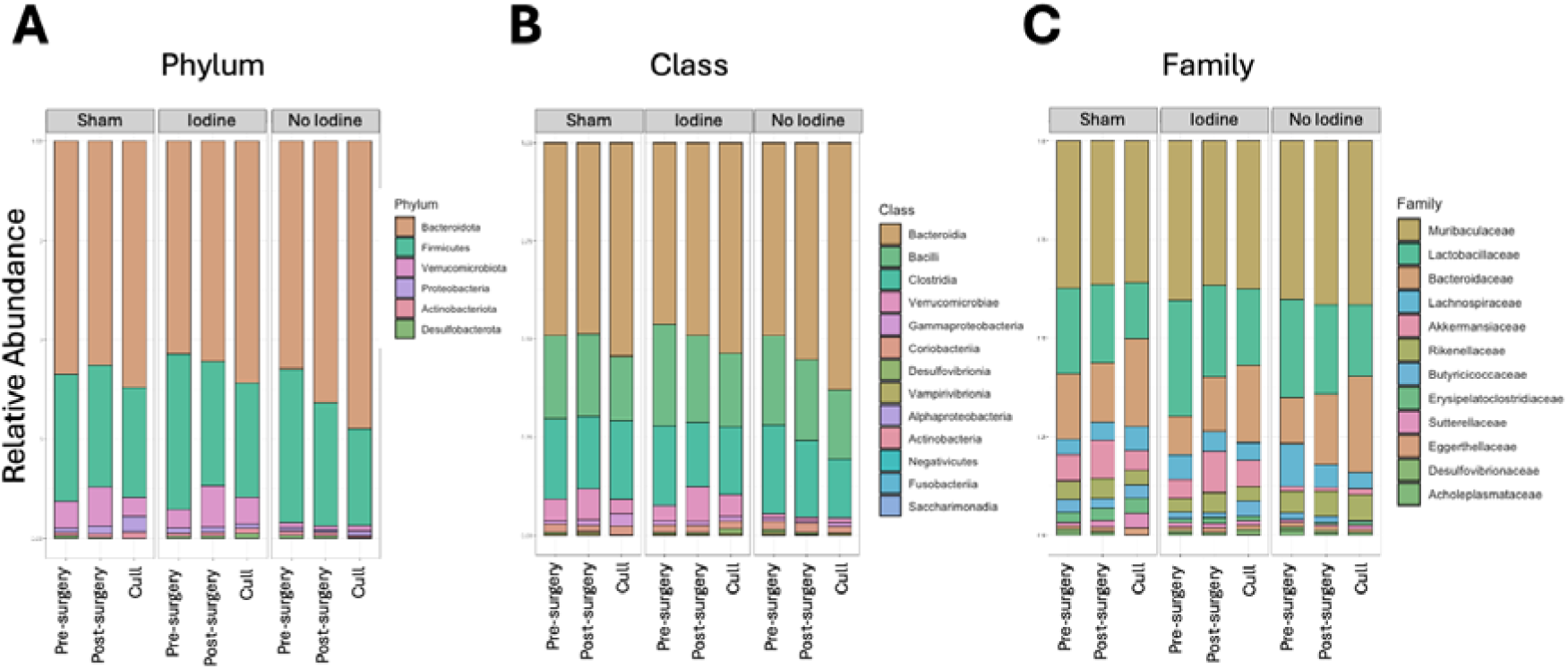
Relative abundance of microbial taxa pre- and post-surgery and at time of cull. Relative abundance of microbial taxa at the **A**: phylum, **B**: class, and **C:** family level. Each bar is color-coded to denote the proportion of the respective phyla, classes or family present. These plots provide a comparative overview of the microbial community structure across pre-surgery, post-surgery, and cull phases of the study in samples from mice receiving sham, non-iodinated (“no-iodine”) mesh and ICP mesh implants. Control: sham, no-iodine: non-iodinated mesh, ICP mesh: iodine.

Muribaculaceae was the most abundant Family and remained prevalently abundant across all phases, driving the observed predominance of phylum Bacteroidota **(Figure 5)**. An overall trend toward increased Bacteroidaceae at the time of culling was also observed in all groups. A marginal increase in the proportion of Gammaproteobacteria was observed in the sham control group over time, which was not seen in the ICP or non-iodinated mesh treatment groups.

Interestingly, Akkermansiaceae (phylum Verrucomicrobiota), a family known for its critical role in gut health and function, was significantly less abundant in the no-iodine mesh group compared to the sham and iodine mesh groups. This substantial shift in baseline microbiome composition persisted across all time points in the no-iodine mesh group. In contrast, Akkermansiaceae was the fourth most abundant family in the sham and iodine mesh groups and exhibited an increase in relative abundance post-surgery. Additionally, the no-iodine mesh group displayed notable microbiome fluctuations over time that were not observed in the sham and iodine mesh groups. These fluctuations included a decline in the abundance of Lachnospiraceae and Butyricicoccaceae, both of which include genera known to produce key gut metabolites such as propionate and butyrate ^46^.

The observation that one group of mice at baseline (i.e., samples taken prior to surgery from mice allocated to the non-iodinated mesh implantation group) were lacking in Akkermansiaceae is likely due to these mice being housed together in a single cage during transportation to the animal facility. This “cage effect” in which differences in the microbiome develop due to the segregated housing environment is well established in the literature. To control for this environmental impact, subsequent studies will ensure that mice from individual transportation cage environments (and cage-specific microbial profiles) are cohoused with mice from other cages when allocated to treatment groups.

#### Alpha diversity

Species richness, as measured by the Chao1 index, showed no statistically significant differences between the pre- and post-surgery phases within the control (sham) and non-iodinated mesh groups (**Figure 6A**). In contrast, the iodine mesh group exhibited a statistically significant decrease in species richness between the pre- and post-surgery phases (p = 0.048) and between the pre-surgery phase and the time of cull (p = 0.045). While these results suggest a potential loss of richness in the iodine mesh group, a similar, albeit non-significant, trend was observed in the sham group, which underwent a mock surgical procedure. This indicates that the observed changes may be subtle and require increased statistical power to confirm in other groups. Moreover, We observed no statistically significant differences in species evenness across groups or phases (**Figure 6B**), indicating relatively consistent community composition and distribution within each group throughout the study.

**Figure 6.**
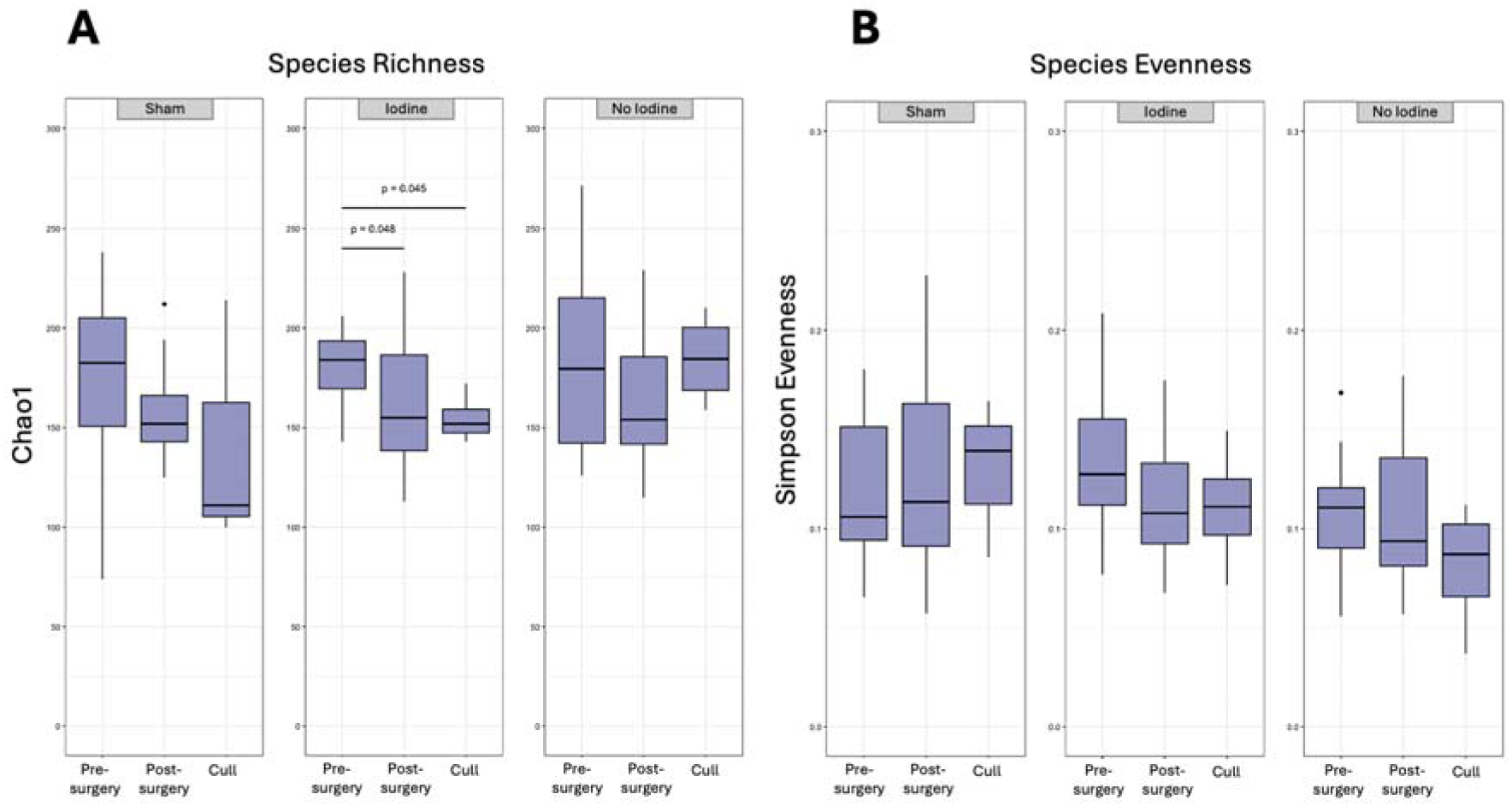
Alpha diversity analysis of species evenness and richness within the gut microbial community. **A:** Species richness and **B:** evenness for control (sham), iodine mesh, and no-iodine mesh groups at pre-surgery, post-surgery, and at time of cull.

Overall, the stable species richness in the control and non-iodinated mesh groups suggests minimal disruption to microbial communities during the experiment.

Analysis of Shannon diversity, which reflects both species richness and evenness, showed no significant differences across phases in the control (sham) and non-iodinated mesh groups (i.e., pre-surgery, post-surgery, and at the time of cull). In contrast, the iodine mesh group exhibited a decline in Shannon diversity at the post-surgery timepoint, primarily driven by reduced richness relative to the pre-surgery baseline (**Figure 7**). However, post-surgery diversity in the iodine mesh group was comparable to that of the sham and non-iodinated mesh groups. Overall, the observed changes across all groups are marginal and do not provide strong evidence for significant disruptions to gut microbial diversity.

**Figure 7.**
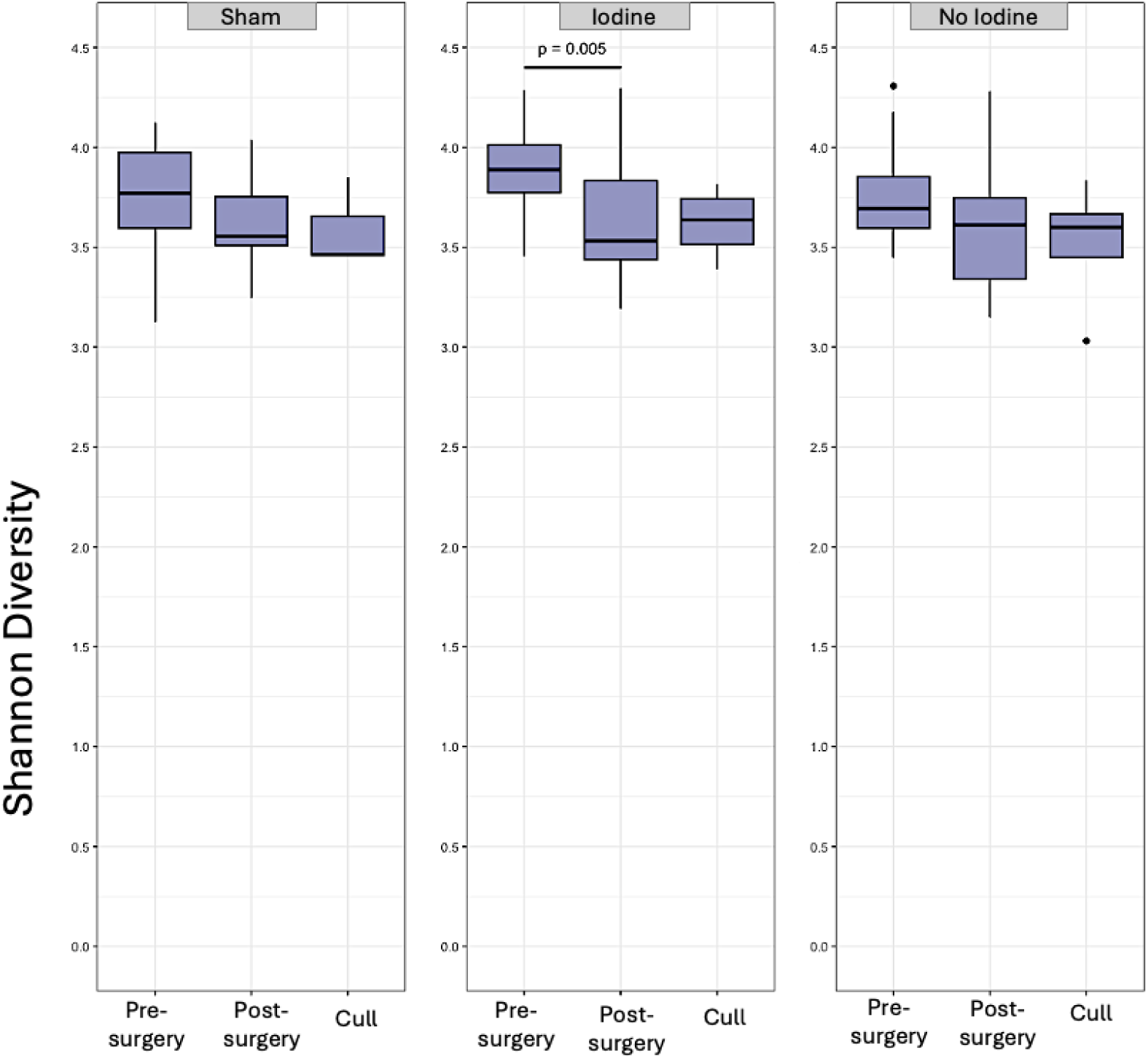
Alpha diversity analysis illustrating the overall diversity of gut microbial community samples for control (sham), iodine mesh, and no-iodine mesh groups at pre- surgery, post-surgery, and at the time of cull.

#### Beta diversity

Beta diversity analysis comparing the sham control to iodinated and non-iodinated mesh groups pre- and post-surgery revealed statistically significant differences among the four groups, driven primarily by compositional changes between pre- and post-surgery phases (p = 0.046 for sham versus iodinated mesh; p = 0.040 for sham versus non-iodinated mesh). However, post-hoc pairwise comparisons did not identify any statistically significant differences, likely due to limited statistical power when comparing smaller groups and the subtle nature of compositional changes, as suggested by the p-values being near the significance threshold (i.e., ≥ 0.04).

To illustrate non-significant trends in compositional differences between the sham and mesh groups, ordinations are presented for pre- and post-surgery groups separately (**Figure 8**). Post-surgery, the microbial communities of the sham control and iodinated mesh groups appear more similar to each other (**Figure 8A**), potentially reflecting the shared increase in Akkermansiaceae as shown by relative abundance data at the family level. In contrast, the sham control and non-iodinated mesh groups exhibited distinct compositions pre-surgery, a pattern that persisted post-surgery, likely due to the unique baseline microbiome composition of the non-iodinated mesh group (also demonstrated in relative abundance analyses). Given these baseline differences, interpreting post-surgery compositional changes is challenging, and the observed trends in ordination should be viewed as exploratory rather than conclusive.

**Figure 8.**
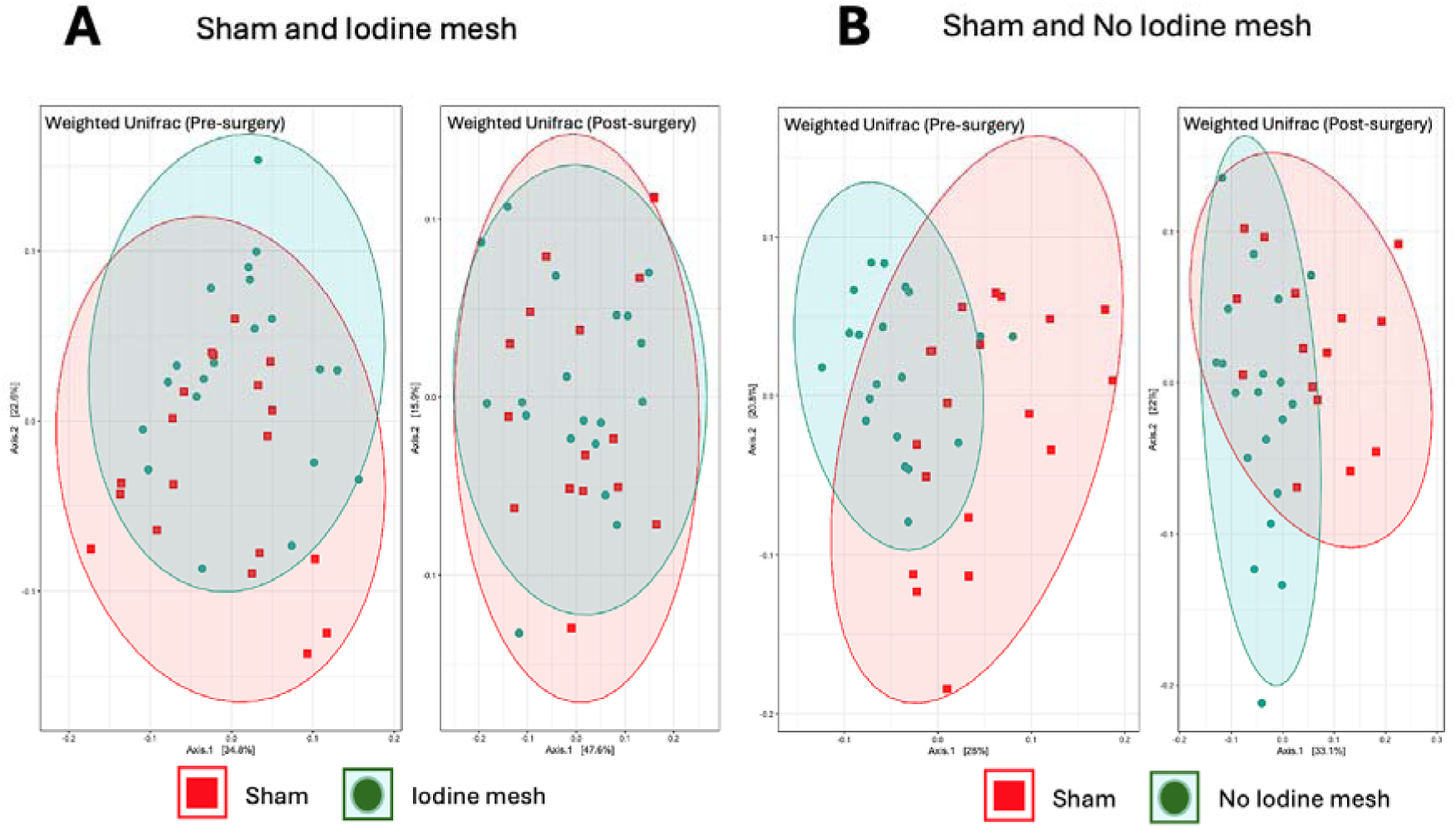
Principal coordinate analyses visualising (dis)similarity of microbial community compositions. **A**: Control and iodinated mesh groups pre-surgery (left) and post-surgery (right). **B.** Control and non-iodinated mesh groups at pre-surgery (left) and post-surgery (right). Each data point represents an individual fecal sample; red: sham control, and green: mesh.

## Conclusion

We report the development of a novel, visible, and biocompatible polymeric mesh aimed at overcoming the limitations associated with traditional transvaginal implants for PFDs. The composite mesh, composed of polymethylmethacrylate (PMMA), and thermoplastic polyurethane (TPU), integrates iodinated carbon nanoparticles (ICPs) to enhance visibility through CT imaging and is coated with 2-methacryloyloxyethyl phosphorylcholine polymer (PMPC) to minimise protein adsorption and promote tissue regeneration. The mechanical properties of the mesh closely resemble those of the native vaginal wall, ensuring a balance of flexibility and strength for optimal performance in supporting prolapsed organs while significantly enhancing visibility through CT imaging.

Results from both cell culture and *in vivo* studies support the potential for the use of this mesh in surgical applications. *In vitro* studies demonstrated high cell viability and reduced protein adsorption, confirming the excellent biocompatibility of the mesh. The addition of PMPC coating further reduced protein adhesion, a key factor in minimising inflammatory responses. *In vivo* evaluations in mice revealed no overall impact on animal health and no change in spleen weight with mesh implantation suggesting that potential inflammatory responses were well tolerated. Nevertheless, we observed increased levels of some cytokines.

Analyses of fecal microbial samples revealed that implantation of the iodine mesh did not significantly affect microbiome composition. The primary shifts in taxonomic composition were an increase in Akkermansiaceae, observed post-surgery in both the sham and iodine mesh groups and the absence of Akkermansiaceae in the baseline community of the non-iodinated mesh group, likely resulting from cage effects during transportation and housing.

We observed a slight decline in species richness and Shannon diversity post-surgery in the iodine mesh group; however, a similar non-significant trend was noted in the sham group, suggesting that higher statistical power might reveal this effect to be related to surgery rather than the iodine mesh itself. Overall, our findings indicate minimal evidence that the iodine mesh specifically impacted the microbiome

To reduce foreign body responses to implantation and improve healing via the addition of secreted growth factors, future research will incorporate the use of patient-rich plasma technology (PRP) ^47^ in which mesh implants will be coated with recipient platelets. Further research is also needed to investigate histological measures of inflammation to determine if increased cytokine levels correspond with structural changes in tissues adjacent to the surgical implant site. Future studies should also explore more extended recovery periods to determine whether mice receiving sham surgery or mesh implants (with or without iodinated ICPs) exhibit similar recovery patterns and cytokine profiles at later time points post-surgery.

Overall, our findings suggest that the composite mesh has the potential for safe implantation, in addition to offering biocompatibility, mechanical compatibility, and novel imaging properties. Further research is required, however, to explore whether adverse immune responses can occur and to characterise recovery profiles over an extended period. Further modifications such as the inclusion of platelet-rich plasma (PRP) technology utilising the recipient’s own blood will be implemented to enhance tissue healing and reduce immune responses following mesh implantation. The innovative mesh reported here offers significant promise for reducing complications associated with existing implants while enabling a novel approach for non-invasive post-surgical monitoring. Future clinical trials will be essential to validate its potential and efficacy as a viable solution for treating PFD, ultimately improving long-term patient care.

