## Supplementary Info for "Bioengineered visible polymeric mesh to enhance urogynaecological health"

### Supporting information

#### XRD and FTIR graphs

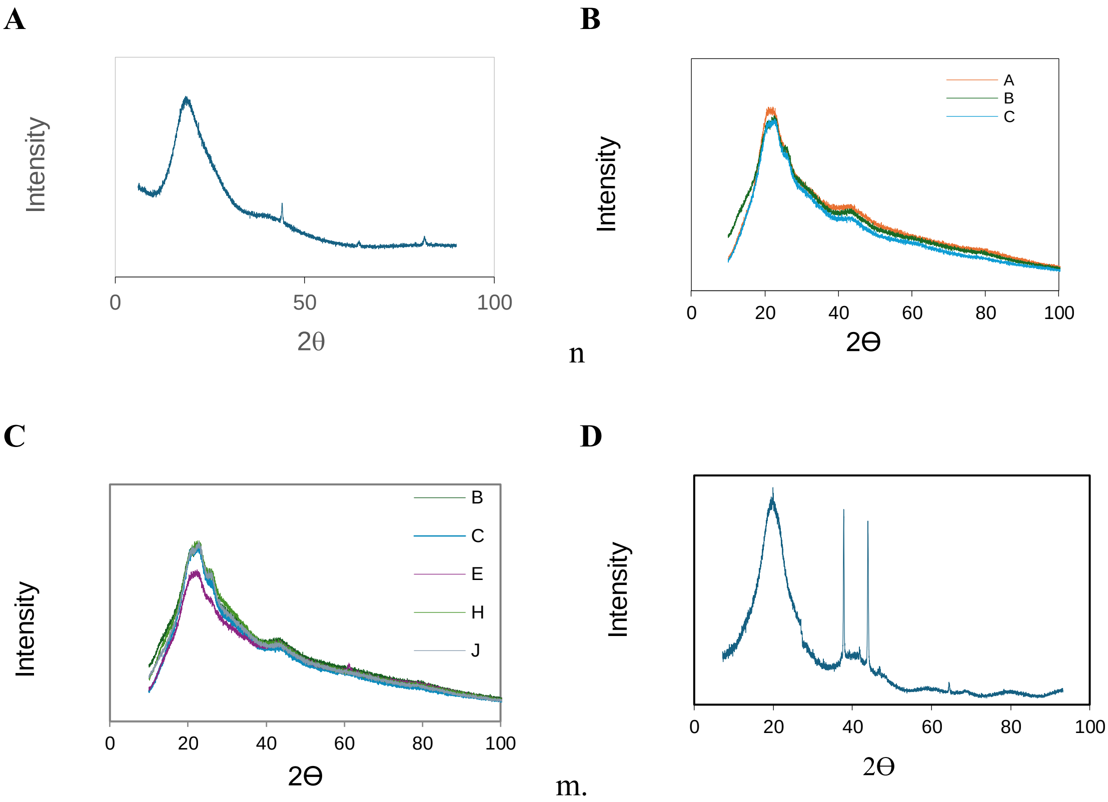


**Supplementary Figure 1.** XRD spectra of (A) ICPs (B-C) filament with various ICPs concentrations (D) 70% ICPs recorded with the second method explained in the manuscript.

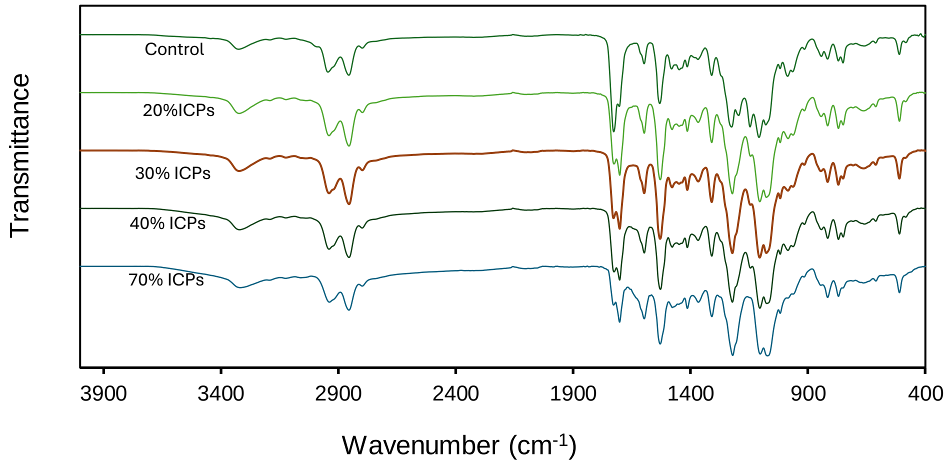


**Supplementary Figure 2.** FTIR spectra of the PMMA/TPU (2:3) filaments with various ICPs concentrations 0%, 20%, 30%, 40% and 70 wt. % of PMMA/TPU in the core section of the filament.
